# Diverse roles of guanine nucleotide exchange factors in regulating collective cell migration

**DOI:** 10.1101/076125

**Authors:** Assaf Zaritsky, Yun-Yu Tseng, M. Angeles Rabadan, Shefali Krishna, Michael Overholtzer, Gaudenz Danuser, Alan Hall

## Abstract

Efficient collective migration depends on a balance between contractility and cytoskeletal rearrangements, adhesion, and mechanical cell-cell communication, all controlled by GTPases of the RHO family. By comprehensive screening of guanine nucleotide exchange factors (GEFs) in human bronchial epithelial cell monolayers, we identified GEFs that are required for collective migration at large, such as SOS1 and *β*-PIX, and RHOA GEFs that are implicated in intercellular communication. Downregulation of the latter GEFs differentially enhanced front-to-back propagation of guidance cues through the monolayer, and was mirrored by downregulation of RHOA expression and myosin-II activity. Phenotype-based clustering of knock-down behaviors identified RHOA-ARHGEF18 and ARHGEF3-ARHGEF28-ARHGEF11 clusters, indicating that the latter may signal through other RHO-family GTPases. Indeed, knock-down of RHOC produced an intermediate between the two phenotypes. We conclude that for effective collective migration the RHOA-GEFs-→ARHOA/C→ actomyosin pathways must be optimally tuned to compromise between generation of motility forces and restriction of intercellular communication.

## Introduction

Collective cell migration involves intercellular mechanical communication through adhesive contacts (Tambe et al., 2011; Weber et al., 2012; Zaritsky et al., 2015). In migrating monolayers such communication is initiated by cells at the monolayer boundary (aka, *leader cells)* and gradually transmitted to cells at the back of the group (Ladoux et al., 2016; Mayor and Etienne-Manneville, 2016; Ng et al., 2012; Serra-Picamal et al., 2012; Zaritsky et al., 2014; Zaritsky et al., 2015). Effective cell-cell communication requires balanced control of contractility, cell-cell and cell-matrix adhesions(Bazellières et al., 2015; Cai et al., 2014; Das et al., 2015; Hayer et al., 2016; Hidalgo-Carcedo et al., 2011; Notbohm et al., 2016; Plutoni et al., 2016; Weber et al., 2012). Coordination between these processes is regulated, among several pathways, by signaling activities of the Rho family GTPases (Cai et al., 2014; Hidalgo-Carcedo et al., 2011; Omelchenko and Hall, 2012; Omelchenko et al., 2014; Plutoni et al., 2016; Reffay et al., 2014; Timpson et al., 2011; Wang et al., 2010). Rho family GTPases are spatially and temporally modulated by complex networks of upstream regulators, including 81 activating guanine nucleotide exchange factors (GEFs), 67 deactivating GAPs, and 3 guanine dissociation inhibitors (GDIs) (Jaffe and Hall, 2005; Omelchenko and Hall, 2012). The networks are composed of many-to-one and one-to-many interaction motifs, i.e., individual GTPases are regulated by multiple GEFs and one GEF often acts upon multiple GTPases. Moreover, some GEFs are effectors of GTPases, leading to nested feedback and feedforward interactions (Cherfils and Zeghouf, 2013; Hodge and Ridley, 2016; Jaffe and Hall, 2005; Schmidt and Hall, 2002). Such pathway ‘design’ permits an enormous functional specialization of transient signaling events, at specific subcellular locations and with precise kinetics.

Our long term goal is to disentangle these signaling cascades in the context of collective cell migration. Although the roles of GEFs and their interactions with Rho GTPases are widely studied for single cell migration (Goicoechea et al., 2014; Pascual-Vargas et al., 2017), less is known how they regulate collective migration (Hidalgo-Carcedo et al., 2011; Omelchenko et al., 2014; Plutoni et al., 2016). Here, we report a comprehensive and validated, image-based GEF screen that identified differential roles of GEFs. By design of quantitative measures that encode the collective dynamics in space and time we were able to identify a surprising role of RHOA, RHOC and a group of four upstream GEFs in modulating collective migration via efficient long-range communication

## Results and Discussion

### Quantification of monolayer cell migration in space and time

Collective cell migration emerges from the individual motility of cells in an interacting group: An action of one cell affects its neighbor and can propagate over time to eventually coordinate distant cells (Zaritsky et al., 2015). To identify molecules implicated in this mechanism we performed live cell imaging of the wound healing response of human bronchial epithelial cells from the 16HBE14o (16HBE) line (Fig. 1A, Video S1). Cells formed apical junctions and maintained epithelial markers and group cohesiveness before scratching the monolayer, as assessed by the localization of E-Cadherin and the tight-junction protein ZO1 at the lateral cell--cell contact areas (Fig. 1B). Upon scratching, the monolayer transitioned over ~2 hours from a non-motile phase to an acceleration phase to steady state wound closure (Fig. 1C). The acceleration phase was associated with a gradual transition of cells from unorganized local movements to a faster and more organized motility. Cells at the wound edge underwent this transition first, followed by a wave of coordinated motility propagating away from the wound edge (Fig. 1A, insets). The propagation is thought to be driven by mechanical cell-cell communication (Matsubayashi et al., 2011; Ng et al., 2012; Notbohm et al., 2016; Serra-Picamal et al., 2012; Zaritsky et al., 2014; Zaritsky et al., 2015).

**Figure 1:**
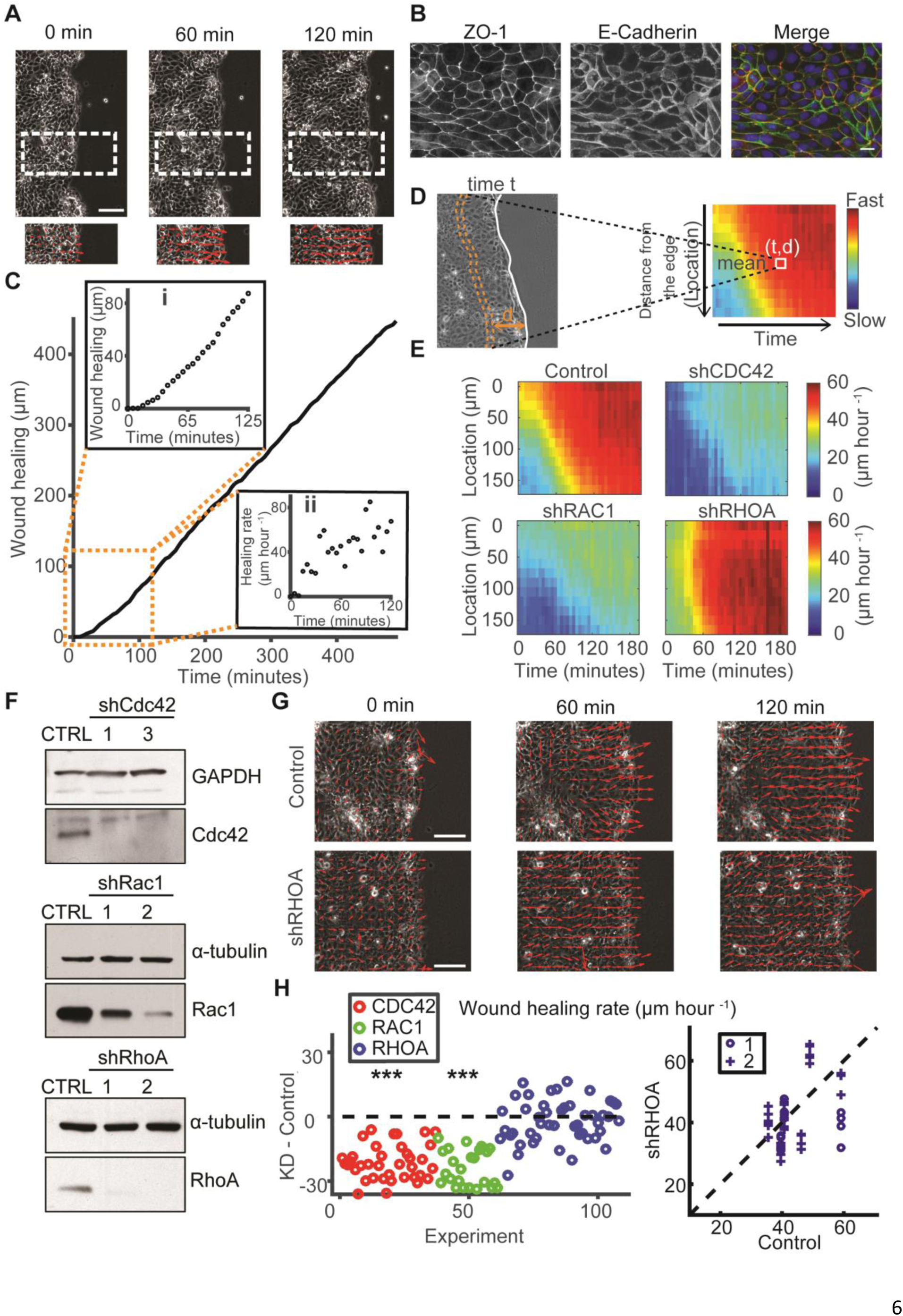
Effects of GTPase knockdown on collective cell migration. (A) Example of a wound healing experiment. Insets: velocity vectors showing that front cells begin to migrate before deeper cells. Scale bar = 100 *μ*m. (B) Immunofluorescent staining of E-Cadherin and ZO1 before scratching the monolayer. Scale bar = 20 *μ*m. (C) Monolayer edge evolution over 500 minutes. Inset top: edge evolution during the first 125 minutes. Inset bottom: increase in of wound healing rate during the first 125 minutes. (D) Construction of a kymograph of a wound healing experiment – mean speed of cells at different distances from the monolayer edge over time. (E) Speed kymographs for control cells and under depletion of Cdc42, Rac1, and RhoA. All kymographs are averages of 4 locations in a well (similar to (Kim et al. 2013)). (F) Western blots of Control, RhoA-, Rac1-, and Cdc42-depleted cells. (G) Monolayer velocity over time (left-to-write) for control (top) and RhoA-depleted cells (bottom). Scale bar = 100 μm. (H) Wound healing rate. Each point was calculated as the difference between the migration rate in one location and the mean of the same day’s control experiment. N = 6-9 days, n = 24-47 locations (CDC42: N = 9, n = 37; RAC1: N = 6, n = 24; RHOA: N = 9, n = 47). Statistics via Wilcoxon signed rank test: p ≤ 0.01: *, p ≤ 0.001: **, p < 0.0001: ***. The right panel shows the migration rate in one location as a function of the mean daily control for the two RhoA hairpins. Such hairpin-specific visualizations of other experiments presented throughout this manuscript are available online (see Methods, Data availability).

To screen for GEFs implicated in the migration-initiating process, we devised an automated and robust pipeline quantifying the spatiotemporal dynamics of motility activation in the monolayer. The core of this pipeline relies on robust detection of the wound edge via a segmentation algorithm that considers both image texture and segmentation consistency over time (Methods). We then applied particle image velocimetry at sub-cellular resolution to measure the speed of cells in the monolayer as a function of their distance to the edge over time. Spatiotemporal dynamics were quantified and visualized in kymographs, in which every bin records the mean speed in a band of constant distance from the wound edge at a particular time point (Fig. 1D). As expected, the speed kymograph showed a backward propagating wave (Fig. 1E, top-left).

Using this assay we first measured the effects of depletion of the canonical Rho GTPases Cdc42, Rac1 and RhoA. Short hairpin RNAs (shRNAs) were designed to selectively knockdown each GTPase (Fig. 1F). As expected (Omelchenko and Hall, 2012; Plutoni et al., 2016; Simpson et al., 2008; Vitorino and Meyer, 2008), knockdown of Cdc42 and Rac1 reduced cell speeds throughout space and time (Fig. 1E, Fig. 1H - wound healing rate; confirmed by a second distinct hairpin Fig. 1F, Fig. S1A). We also confirmed inhibition of cell speed and monolayer migration under shRNA-mediated knockdown of the Rac1- and Cdc42-GEF *β*-PIX (Omelchenko et al., 2014) (Fig. S1B-D).

Knockdown of the Rho-GTPase RhoA induced near concurrent transition of cells at the front and back of the monolayer from a non-motile to a motile state (Figs. 1E, 1G). Importantly, RhoA-depleted cell monolayers reached the same wound healing rates as unperturbed monolayers (Fig. 1H). These data suggested that RhoA is critically implicated in the communication chain required to synchronize cell behavior throughout the monolayer after wound infliction.

### Information encoded in a wound healing experiment

Given the differential effects observed for the knockdown of Rho GTPases we expected that down-regulating GEFs might also differentially alter the dynamics of motility activation. To capture the shifts in these behaviors we established a concise representation of a live wound healing experiment between wound infliction and steady state wound closure, which could be compared across many experiments. We defined a 12-dimentional feature vector to encode the information contained in a speed kymograph over the first 180 μm of the monolayer and 200 minutes post scratching by averaging the speed values in bins of 60 μm and 50 minutes, respectively (Fig. 2A, Methods). Accordingly, features 1-4 encode the acceleration of cells at the monolayer front, features 5-8 the acceleration of cells 60-120 μm behind the monolayer edge, etc. Features 1, 5 and 9 encode spatial variations in migration shortly after wounding. In the example of Fig. 2A, cells at front have already developed motility (feature 1), but cells further back in the monolayer (feature 9) have not. Features 2, 6 and 10; 3,7, and 11; and 4, 8, and 12 encode the same spatial variations at later times. The pattern of 12-dimensional feature vectors was replicated for 402 control experiments without shRNA treatment (Fig. 2B, top). The broad variation in collective migration behavior, even for constant experimental conditions, was visualized by normalizing each feature across the population of all 402 experiments (Fig. 2B, bottom; see Methods).

To identify core features that would capture the relevant variation of these data over noise, we applied *Principal Component Analysis* (PCA) (Jolliffe, 2002) to orthogonally-transform the set of possibly correlated 12-dimensional features to a smaller set of linearly-uncorrelated features denoted as the *principal components* (PCs). Indeed, when using PCs, the first 3 components captured 92% of the variability in the population of control experiments, allowing a four-fold dimensionality reduction of the feature space. The variation of PC1 between experiments was highly correlated with the variation in wound healing rate (Fig. 2C). The same applied to a lesser extent to PC3, whereas PC2 was uncorrelated. We further analyzed the coefficients that map the original 12 features into the scalar value of the respective PC. For PC1 all 12 coefficients have a similar value. Hence the mapping corresponds to an averaging of the speed values across the kymograph (Fig. 2D). For PC2, the coefficients are positive for features 1, 5, 9, i.e., the speed values in the first time window, and negative for features 4, 8, 12, i.e., the speed values in the last time window. Thus, PC2 captures the reverse temporal gradient of the speed values. For PC3, the coefficients are positive for features 1 - 4, i.e., the speed values at the wound edge and negative for the features 9 - 12, i.e., the speed values in the back of the monolayer. Thus, PC3 captures the spatial gradient of the speed values in the direction of wound closure.

**Figure 2:**
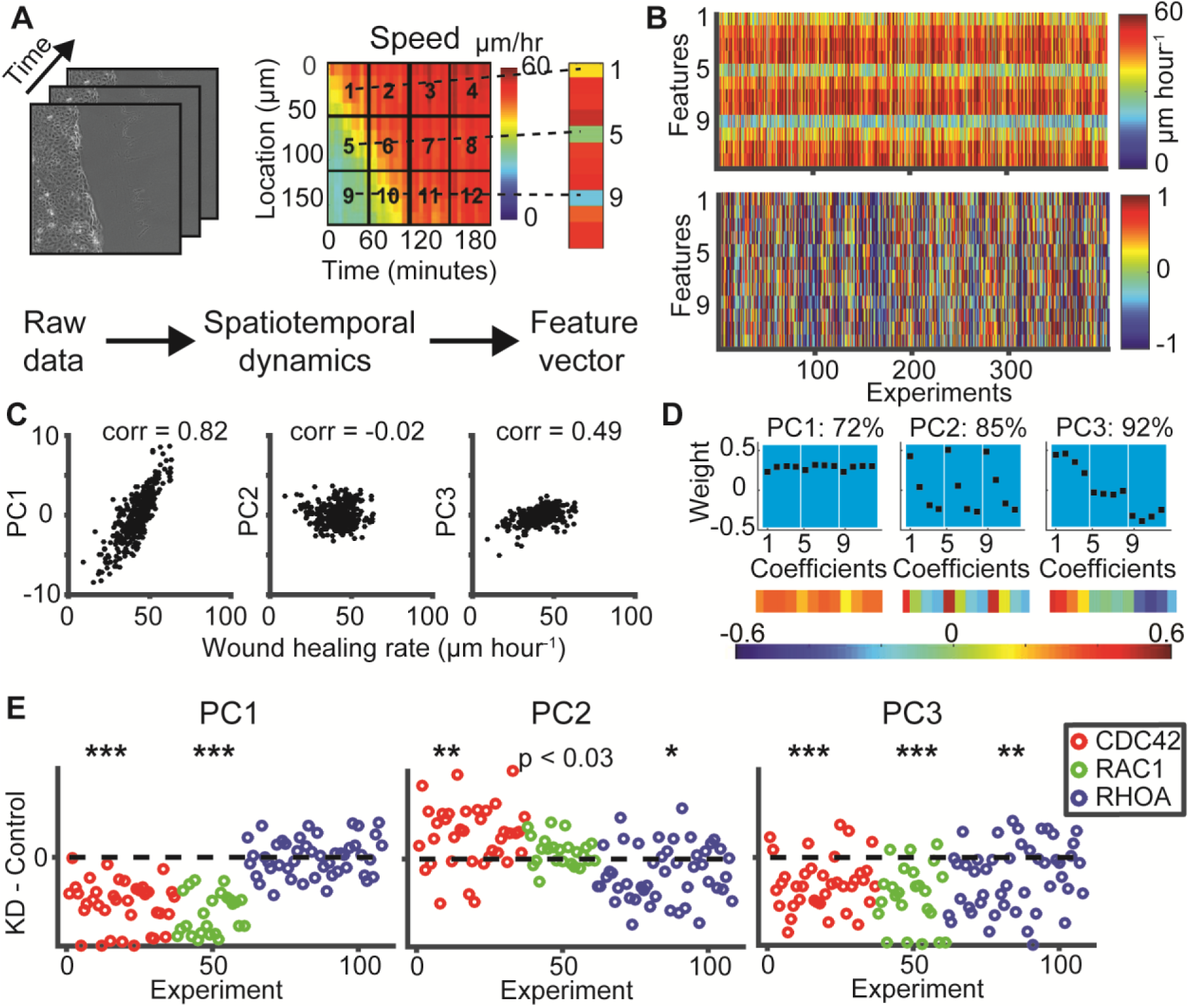
Information encoded in a wound healing experiment. (A) Reducing a wound healing experiment to a 12-dimensional feature vector encoding spatiotemporal dynamics. (B) Speed feature vectors for 402 control experiments. Bottom panel displays the normalized features. (C) Association of Principal Components (PCs) 1-3 with wound healing rate. (D) Coefficients of PC 1-3, together encoding 92% of the speed variance in control experiments. (E) Effect on PCs by depletion of Cdc42, Rac1 and RhoA. Statistics via Wilcoxon signed rank test: p ≤ 0.01: *, p ≤ 0.001: **, p ≤ 0.0001: ***.

Knockdown of Cdc42 and Rac1 decreased PC1 reflecting a significant overall reduction in speed (Fig. 2E). PC2 was somewhat increased for shCdc42 but unaltered for shRac1, whereas PC3 strongly decreased for both. This shows that the global reduction in speed under these conditions is accompanied by flatter spatiotemporal gradients (Fig. 2E, Methods). Knockdown of RhoA most prominently altered PC3 with no effect on PC1 (Fig. 2E), reflecting an alteration in the spatial gradient in accordance with Fig. 1E.

When a monolayer migrates collectively, cells within the monolayer eventually have to migrate toward the monolayer edge. This directional cue is transmitted via backward propagation of a strain gradient (Zaritsky et al., 2014). Thus, the spatial propagation of migration directionality is a measure related to intercellular communication. To capture this aspect we defined for every kymograph bin directionality as the absolute ratio between the mean velocity component perpendicular to the monolayer edge and the velocity component parallel to the monolayer edge. The directionality kymograph displayed a backward propagating wave pattern as previously observed for other cell systems (Ng et al., 2012; Zaritsky et al., 2014) (Fig. S1E). Analogous to the analysis of the speed kymographs, we binned directionality kymographs into a feature vector of 12 values (Methods, and Fig. 2A) and applied PCA. The first three PCs resembled the PCs for speed (Fig. S1F), suggesting a functional coupling between speed and directionality establishment in the process of activating collective migration. Similar to the speed analysis, PC1 of the directionality correlated strongly with the wound healing rate, however, PC2 or PC3 were uncorrelated (Fig. S1G). Upon knockdown of the GTPases, Cdc42 and Rac1 depletion primarily affected the overall magnitude of directionality, whereas RhoA depletion reduced the spatial gradient, indicative of rapid long-range communication between monolayer front and back after wound infliction (Fig. S1H). Altogether, our data established the first three PCs of a feature space derived from kymographs as measures for the detection of alterations in magnitude, temporal or spatial gradients in speed and directionality and underline the roles RhoA plays in long-range communication across the monolayer.

### Comprehensive GEF screen

Equipped with our image-based assay for collective migration we targeted the 81 Rho family GEFs known in the human genome by small hairpin RNA (shRNA, Fig. 3A). A custom library of pSUPER shRNA retroviruses (3 hairpins per GEF) was used for the screening, targeting 80 out of 81 GEFs. 75 out of the remaining 80 GEFs were confirmed to be expressed in the 16HBE cell line by western blotting or qRT-PCR in cases where no reliable antibody was available (Methods). Western blots/qRT-PCR were used to evaluate the knockdown efficiency of every hairpin and a cutoff of 50% depletion was selected for analysis of wound healing experiments (Fig. 3B). 16 GEFs had knockdown efficiency of less than 50% for all 3 hairpins and thus were eliminated from the screen. Within the remaining group of 59 GEFs the knockdown of 11 GEFs could not be validated by qRT-PCR due to failures in the production of efficient primers. Nonetheless, we included those genes in our imaging-based screen and labeled them as ‘unknown knockdown efficiency’. A total of over 2600 time-lapse movies from the screen and the follow-up experiments were analyzed (Methods). Control videos showed broad day-to-day variability in the wound healing rates (Fig. 3C).

To score the alteration of a knockdown well we quantified in multiple image locations the differences to the control well on the same plate. For every location we extracted PCs 1-3 for speed and directionality. For each of the 6 PCs we quantified the separation of target well and control well using three different distance metrics that balance the intra- and inter-experiment variability (Methods). For each combination of PC- and distance metric, we employed 24 experiments with no measureable knockdown (KD efficiency of 0%) as a baseline to calculate z-scores (Methods).

To identify high-confidence hits we estimated an objective z-score threshold by balancing between false-positive and false-negative rates. As *positive controls* we grouped together known motility reducers in Cdc42, Rac1 and *β*-PIX. Experiments with no measurable knockdown of the target were used as *off-target controls.* Their mean and standard deviation were used to calculate z-scores for all experiments. Distributions of z-scores in the speed PC-measures are shown in Fig. 3D. As expected, the positive controls were most discriminative for PC1. Thus, we exploited the known motility reduction of the positive controls together with the corresponding distribution of off-target controls to estimate a threshold appropriate to identify hits (Methods). The threshold was estimated as 9.8. Because the threshold was standardized relative to the off-target controls, it was transferrable to the other PC-measures.

**Figure 3:**
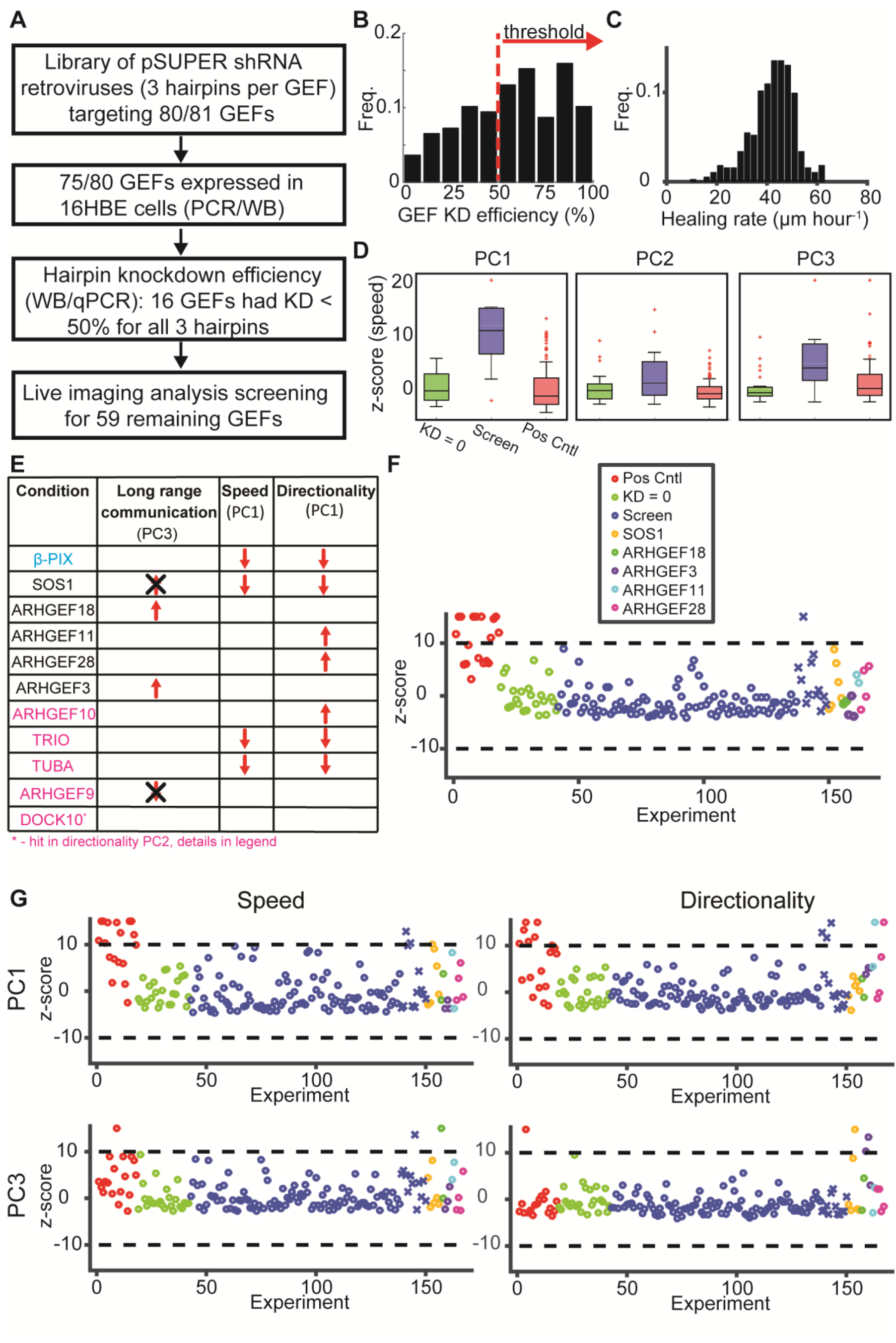
Comprehensive GEF screen. (A) Flowchart of screen. (B) Histogram of knockdown efficiency of all hairpins used in the screen. Only hairpins with depletion greater than 50% where considered for the screen (red). N total number of hairpins = 125. (C) Histogram of wound healing rate. Mean +/− standard deviation: 42.2 +/−8.8 pm hour^-1^. (D) Boxplots of z-scores for off-target controls (green, knockdown efficiency = 0%, N = 24), positive controls (blue, CDC42, RAC1 and *β*-PIX, N = 18) and the screened GEFs (red). Box represents 25-75%, whiskers 5 - 95% of the data, assuming normal distribution. Z-scores were calculated relative to the off-target control. (E) Summary of GEFs identified as hits in the screen and their phenotypes, PC3 in speed or directionality signifies long-range communication. *β*-PIX was a GEF with known effects on collective migration and used as a positive control (cyan). Magenta label hits that were not followed-up. Black cross indicates that these phenoytpes were excluded: see Fig. S2 for details. (F-G) Visualization of measures that were altered by the hits identified in the screen and that were validated. Shown are z-scores for each well. Positive controls (red), off target controls (KD = 0, green), GEFs that were screened (blue) but not identified (circle) or validated (cross) as hits, and validated hits. (F) Wound healing rate. (G) PC1 (top) and PC3 (bottom).

### Screen summary

In addition to *β*-PIX, which has previously been reported as required for collective cell migration (Omelchenko et al., 2014), 10 new GEFs were identified with significant knock-down effects. Among those, we focused on SOS1, ARHGEF18, ARHGEF28, ARHEG11, ARHGEF3 for validation (Fig. 3E, black). The other five (Fig. 3E, magenta, Fig. 3F-G, blue crosses) were not followed-up for reasons discussed with Fig. S2A-E.

### SOS1-RAS pathway is required for monolayer migration

We first investigated SOS1, a dual GEF for RAC1 and RAS (Nimnual et al., 1998) that was found to regulate epithelial tight junction formation in 16HBE cells through the MEK/ERK pathway (Durgan et al., 2014). Knockdown of Sos1 by 2 different hairpins induced a reduction in wound healing rate, cell speed and directionality (Fig. 3F-G, Fig. S2F), consistent with its role as a Rac1 activator. Following the experiments in (Durgan et al., 2014), we then blocked MEK and ERK activity directly by small molecule inhibitors and concluded that the SOS1-RAS pathway (Fig. S2G, Methods) is also required for collective migration, in addition to epithelial tight junction formation (Fig. S2H).

### RhoA GEFs regulate intercellular communication

Four paralog RHOA-GEFs were identified as hits (Fig. 3E). Depletion of these GEFs did not dampen wound healing rate, cell speed, or directionality, but enhanced front-to-back propagation of motility. Specifically, depletion of Arhgef18, similarly to depletion of RhoA, accelerated front-to-rear long-range communication (Fig. 4A). Depletion of Arhgef 3, Arhgef11 and Arhgef28 somewhat accelerated long-range communication in speed and directionality, and enhanced directionality overall (Figs. 3G, 4B-E). GEF-H1 (also called Lcf or ARHGEF2), a mechano-responsive RhoA-specific GEF and paralog of the 4 GEFs identified in the screen (Birkenfeld et al., 2007; Chang et al., 2008; Guilluy et al., 2011b) was excluded from the list because all hairpins produced a knockdown less than 50%. Nonetheless, experiments utilizing these relatively inefficient hairpins revealed a phenotype similar to depletion of RhoA (Fig. 4F, compare to Fig. 2E and S1H for RhoA).

In a set of validation experiments each RHOA-GEF hit was replicated with at least 2 different hairpins in 6-10 wells (total of 23-53 locations per hit). When assessing multiple replicates, systematic alterations were recognized that were missed by the stringent criteria of the screen (Fig. 4G). For example, faster long-range communication in directionality was a common outcome of depleting any RhoA-GEF, while increased directionality (Arhgef11, Arhgef28 and Arhgef3) or long-range communication in speed (Arhgef18) were GEF-specific. This suggests differential roles among RhoA-GEFs in regulating functions down-stream of RhoA signaling.

**Figure 4:**
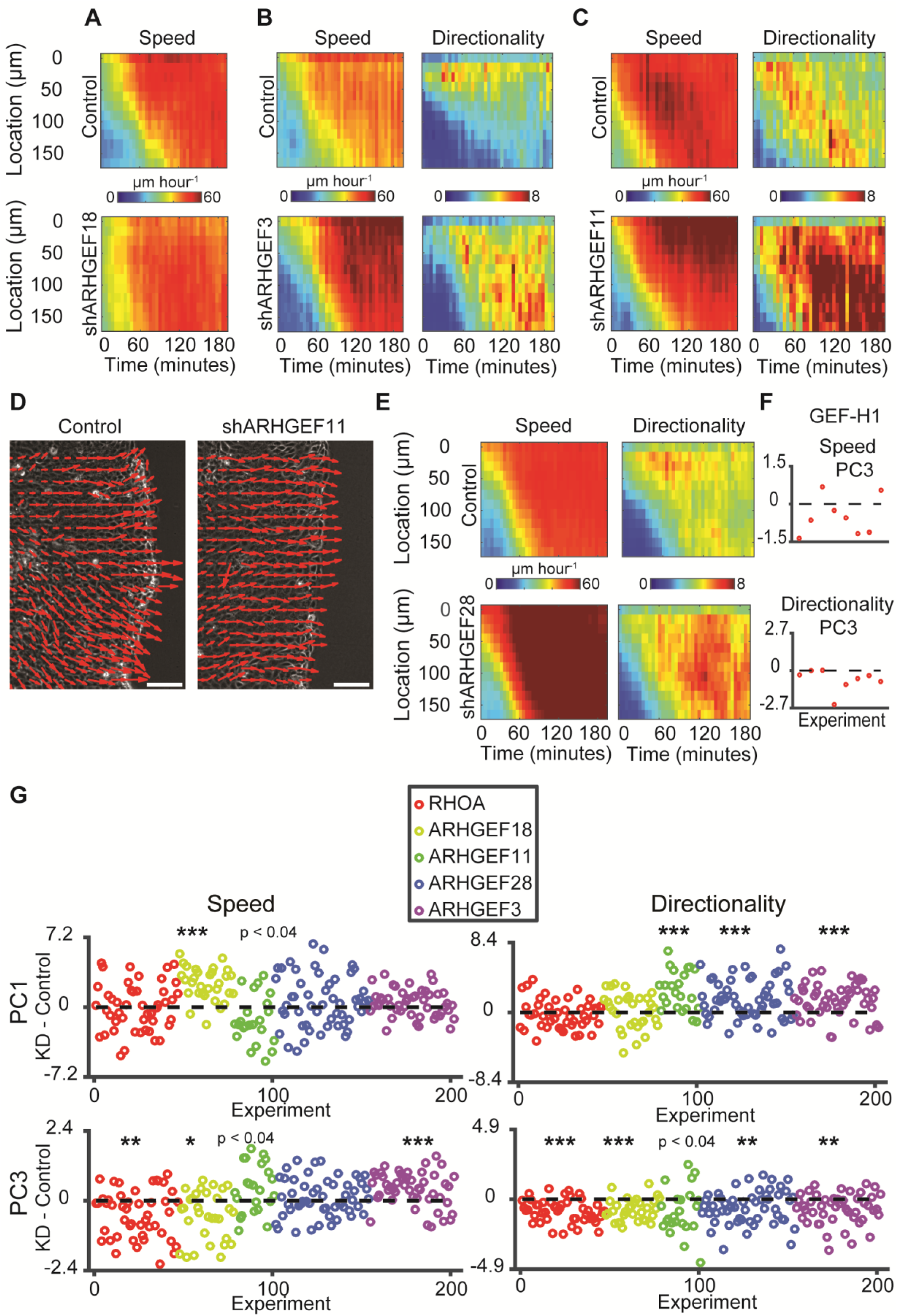
Effects of RhoA GEF depletion on intercellular communication. (A-C) Kymographs of speed and directionality for control vs ARHGEF18, ARHGEF3, and ARHGEF11 knockdown. (D) Vector fields 100 minutes post wound infliction for control vs ARHGEF11 knockdown. Scale bar = 100 μ-m. (E) Kymographs of speed and directionality for control vs ARHGEF28 knockdown. (F) Difference in PC3 for speed (top) and directionality (bottom) between GEF-H1 knockdown (2 hairpins with 30% and 45% depletion) and control. Each point represents the difference in one well location and the mean of the same day’s control experiment. N = 1 day, n = 8 locations. (G) Validation of screen hits in RhoA GEFs. Each point represents the difference in PC of the indicated variable between GEF KD in one well location and the mean of the daily control; N - number of wells, n - number of locations. RHOA: N = 9, n = 47, ARHGEF18: N = 6, n = 31, ARHGEF11: N = 6, n = 23, ARHGEF28: N = 10, n = 53, ARHGEF3: N = 9, n = 48. Statistics via Wilcoxon signed rank test: p ≤ 0.01: *, p ≤ 0.001: **, p ≤ 0.0001: ***.

Transmission of motility guidance cues from cell to cell results in the formation of clusters of cells moving with coordinated trajectories (Zaritsky et al., 2014) (Methods). Cluster formation was quantified by recording the fraction of the monolayer in which cells migrated coordinately.In control experiments, the fraction increased steadily over time (Fig. S3A), due to an expansion of clusters from the front into the monolayer (Fig. S3B-C). In analogy to speed and directionality the spatiotemporal dynamics of cluster formation was captured by 3 PCs, which encoded magnitude, temporal, and spatial gradients (Fig. S3D). Importantly, these PCs explicitly capture the strength and propagation of *short-range communication.* PC1, which was most associated with the wound healing rate (Fig. S3E), was increased upon depletion of Arhgef28, while depletion of RhoA, Arhgef18 or Arhgef28 increased PC3 in coordination (Fig. S3F), showing that under these manipulations front and rear of the monolayer are coordinated rapidly after wounding.

Altogether, these results establish roles for RhoA and four of its activating GEFs in regulation intercellular communication.

### Actomyosin contractility disturbs intercellular communication downstream of the ARHGEF18 - RHOA pathway

Previous work showed that partial down-regulation of actomyosin contractility enhances passive force transmission through cells, whereas high levels of myosin activity ‘scrambles’ mechanical signals (Ng et al., 2015). Hence we speculated that the enhancement of long-range communication induced by RhoA and RhoA-GEFs depletion related to reduced myosin activity under these perturbed conditions. To test this we inhibited Myosin-II directly using Blebbistatin or via ROCK inhibition using Y27632. To assess the effect of short vs. long-term treatment, the drugs were applied without or with 24 or 48 hours pre-incubation to wounding experiment.

Treatment with low dosages of Y27632 (15uM or 20uM) and Blebbistatin (10uM) in general increased cell speed and coordination (Fig. S3G-H). Pre-incubation for 48 hours led to increased long-range communication in directionality and/or increased coordination similar to the behavior observed upon depletion of RhoA/RhoA-GEFs (Fig. S3G-I). Inhibition of the formin pathway, which is also downstream of RhoA (Higashida et al., 2004) (Fig. S3J), with low dosages of SMIFH2 (5uM or 10uM) (Fig. S3J-K), increased speed, directionality and coordination without changing long-range communication (Fig. S3K). Treatment with higher dosages of Y27632 (25uM, for 48 hours) or SMIFH2 (25uM) inhibited overall motility (Fig. S3G, K).

The effect of a 48 hour drug treatment is expected to be biologically more similar to the effect of protein depletion by shRNA than the effects of more acute treatments thus, we next compared the effects of Blebbistatin and Y27632 treatment over 48 hours to the effects of RHOA/RHOA-GEFs depletion (Fig. 5A). We did not consider formin inhibition in this analysis because of the absence of a communication phenotype that could match the RHOA-GEF depletion phenotypes. Similarity analysis was performed by representing each experiment by a 9 dimensional feature vector composed of the normalized PCs #1-3 for speed, directionality and coordination and calculating the similarity between every pair of conditions with the L1 norm. The pair-wise (symmetric) similarity matrix confirmed distinct, but related functionalities for RHOA/RHOA-GEFs/Contractility in control of intercellular communication (Fig. 5B). ARHGEF11, ARHGEF28 or ARHGEF3 fall in a first cluster, ARHGEF18, RHOA, and Blebbistatin treatment in a second cluster. Treatments with 15uM or 20uM Y27632–while similar among each other – differ in their effect from any of the two other phenotypes. This demonstrates the sensitivity of our analysis to distinguish perturbations of the RhoA-myosin pathway from perturbation of Rock, which in part affects the RhoA-myosin axis, but appears to be implicated in additional pathways driving collective migration. More critically, the sensitivity of our assay predicts that ARHGEF18 (and likely also ARHGEF2) regulate specifically RhoA-mediated activation of contractility, while ARHGEF11, ARHGEF28 and ARHGEF3 co-regulate RhoA-independent pathways that do not converge on myosin-II promoted processes. These may include RHOB/C (but not CDC42 or RAC1) for ARHGEF11 (Jaiswal et al., 2011; Rümenapp et al., 1999), RHOC (but not CDC42 or RAC1) for ARHGEF28 (Bravo-Cordero et al., 2011; Bravo-Cordero et al., 2013; van Horck et al., 2001), and RHOB (but not RHOC) for ARHGEF3 (Arthur et al., 2002), although the abundance of RhoC expression in 16HBE is fourfold less of RhoA (Wallace et al., 2011). Acute inhibition of all Rho isoforms (A, B and C) with the small molecule inhibitor Rhosin (Shang et al., 2012), showed a general motility reduction in a dose-dependent manner indicating that Rho isoforms are required for collective migration (Fig. 5C). We also excluded the possibility of a cross-talk between RhoA and RhoC at the expression level. Western blots verified that knockdown of RhoA did not reduce RhoC (Fig. 5D).

To test the prediction of differential regulation of RhoA and RhoC we performed a set of RhoC knockdown experiments. Depletion of RhoC increased cell speed and coordination (PC1), long-range communication in directionality and coordination (PC3) (Fig. 5E). Pair-wise similarity matrix indicated that RhoC had an intermediate phenotype between the RhoA-ARHGEF18 and the cluster of the other RhoA-GEFs (Fig. 5F-G).

Together, these analyses identified ARHGEF18 as the only RHOA-GEF exclusively activating the RHOA isoform (Blomquist et al., 2000; Herder et al., 2013), albeit biochemically it has also been described as a GEF for RAC1 (but not CDC42) (Niu et al., 2003). Arhgef18 activates RhoA at tight junctions, directly interacting with myosin IIA and regulating tight-junction assembly (Durgan et al., 2014; Kim et al., 2015; Terry et al., 2011; Xu et al., 2013). Thus, we interpret the similarity of defects in long-range communication induced by Arhgef18- and RhoA-depletion and direct inhibition of myosin-II contraction as indication that ARHGEF18 locates specifically at the top of an ARHGEF18/RHOA/MYO-II pathway (Fig. 5H). ARHGEF11, ARHGEF28 and ARHGEF3 on the one hand target the RHOA/MYO-II pathway. On the other hand these three GEFs also target migration directionality, which is unaffected by RhoA-mediated signals. The intermediate phenotype of RHOC could be explained by integration of multiple pathways with similar but differential function (Kafri et al., 2009), and/or dynamic interactions between the molecular components that can be regulated in time and space (Guilluy et al., 2011a). The functional similarity between the ARHGEF18 and RHOC phenotypes, notably in cell speed, could be explained by a similar coil domain structure (Cook et al., 2014) predictive of competition, binding or indirect interactions through other proteins of ARHGEF18 with RHOC. How the ARHGEF18/RhoA-, RhoC- and ARHGEF3,11,28- mediated pathways mechanistically differentiate between long-range communication, speed and directionality will require an analysis of the spatiotemporal activation patterns during migration of these upstream GEFs in conjunction with the targeted GTPases, for example by construction of GEF-activation biosensors.

A second limitation of our approach is its inability to identify intercellular-communication phenotypes upon overall reduction of motility, due to inherent reduction of the spatial and temporal gradients in relation to controls (Fig. 2E, PC3). For example, TRIO’s Rho-targeting GEF domain is activated by a non-canonical Notch signaling pathway (Le Gall et al., 2008; Song and Giniger, 2011), known to be important in intercellular communication (Grego-Bessa et al., 2007; Lai, 2004). Based on our findings of a RhoA-axis in the regulation of intercellular communication it would be obvious to hypothesize that this role of TRIO is modulated by its RhoA targeting DHPH2 domains (Bellanger et al., 2000). Unfortunately, this hypothesis could not be tested by knock-down experiments as performed here. Trio knockdown primarily caused an overall reduction of motility, possibly through its activating interaction with Rac1/RhoG mediated by the DHPH1 domain. A differential analysis of domain-specific effects in GEFs with multiple GTPase interactions is beyond the scope of this work.

Consistent with several other reports (Lim et al., 2010; Vicente-Manzanares et al., 2009), our data suggests again that, while necessary for the generation of motility forces, myosin II-mediated contractility acts as an inhibitor for cell-to-cell communication, likely because contractility intercepts passive mechanical force transduction through the cellular cortex (Ng et al., 2015). Hence, during collective migration myosin-II contractility must be adjusted to balance two opposing objectives: Generation of robust motile forces vs. transmission of mechanical guidance cues. Here we show that signaling pathways upstream of myosin II such as the RHOA/ROCK pathway need to be balanced in the same manner.

**Figure 5:**
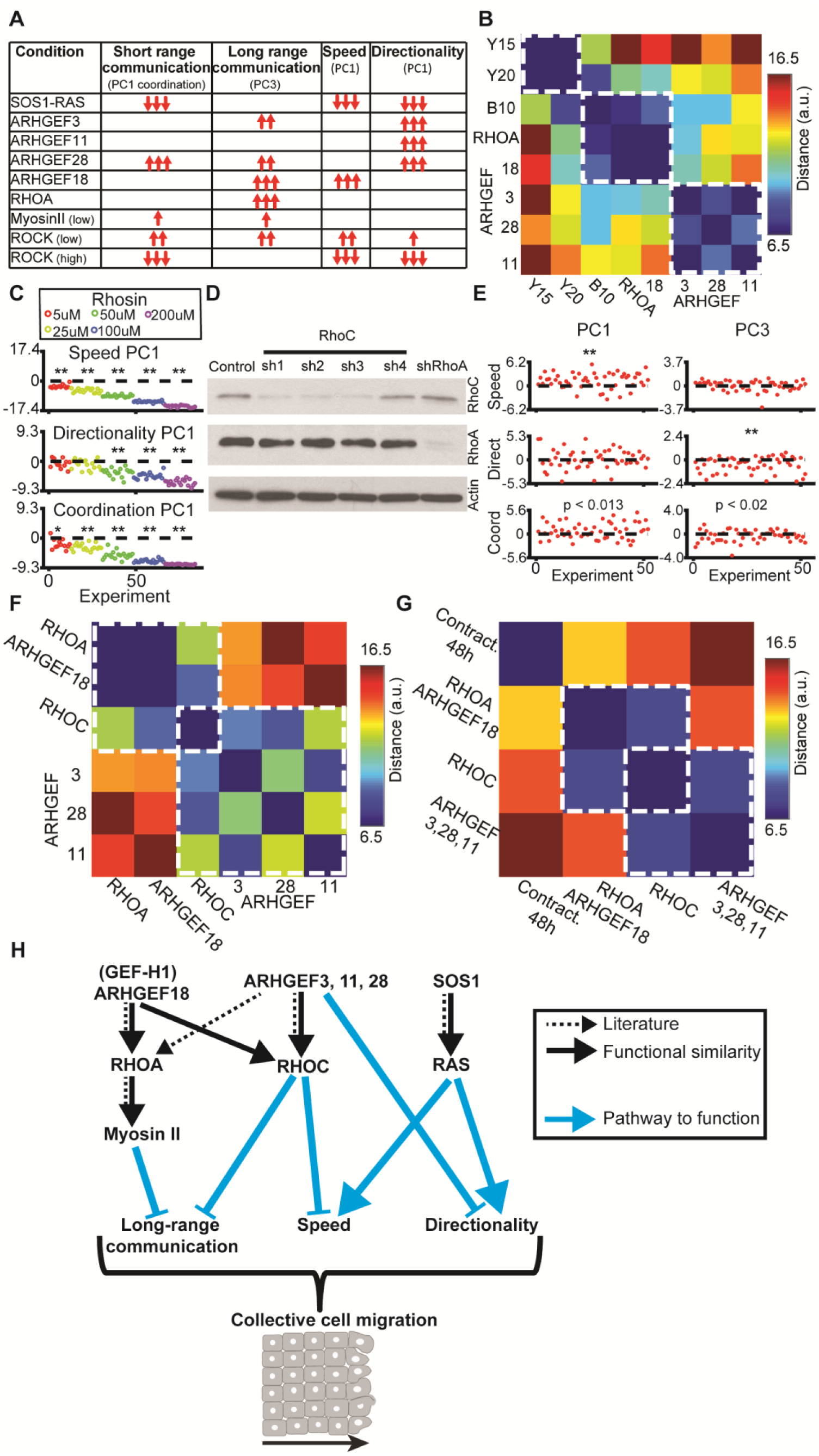
Distinct functional clusters along the RHOA-GEFs/RHOA/actomyosin pathway. (A) Summary of all phenotypic alterations by validated hits of our GEF-screen and for experiments using inhibition of contractility. Number of arrows corresponds to most significant p-value for a given attribute (p ≤ 0.01, p ≤ 0.001 or p ≤ 0.0001 correspondingly). Empty table bins mean ‘no phenotype’. (B) Pairwise distance matrix for knockdown of RhoA, RhoA-GEFs and contractility experiments. Y, Y27632, B, Blebbistatin; number indicates concentration applied over 48 hours prior to wounding. Dashed white boxes show functional clusters as defined by close phenotypic similarity. (C) Treatment with Rhosin (5, 25, 50, 100 or 200 uM) induced motility in a dose-dependent fashion. Each point represents the difference in PC of the indicated variable between drug treatment in one well location and the mean of the daily control; N = 2 and n = 12 for 5uM and N = 3, n = 18 for the other concentrations, 1 control well per daily experiment. Statistics via Wilcoxon signed rank test: p ≤ 0.01: *, p ≤ 0.001: **, p ≤ 0.0001: ***. (D) Western blots to identify expression cross-talk between RhoC- and RhoA-depletion. (E) Effect on PCs by depletion of RhoC. N = 9 and n = 52. Statistics via Wilcoxon signed rank test. (F) Pair-wise distance matrix for knockdown of RhoA, RhoC and RhoA-GEFs. (G) Pair-wise distance matrix the mean clusters phenotype: contractility applied for 48 hours, RhoA and Arhgef18, RhoC, Arhgef3, 28 and 11. (H) Working model emerging from the phenotypic similarity established in this study and existing literature. Dotted black arrows are biochemical associations described in the literature (see citations in text), full black arrows indicate functional similarities and cyan arrows link protein knockdown to specific phenotype.

## Methods

### Cells and culture conditions

The human bronchial epithelial cell line, 16HBE14o- (16HBE), was kindly provided by the laboratory of Dr. Dieter C. Gruenert (University of California, San Francisco). 16HBE cells were cultured in MEM (MSKCC core facility), supplemented with 10% fetal bovine serum (FBS) (Omega Scientific, lot number 169905), GlutaMAX (35050, Gibco), and a mixture of penicillin-streptomycin (100X, 10000 U/mL) (15140, Gibco). Stable cell lines were selected with 1.5 μg/mL puromycin (P7255, Sigma). HEK293T cells were purchased from ATCC, and grown in DME high glucose + sodium pyruvate (MSKCC core facility) supplemented with 10% FBS (Omega Scientific, lot number 169905) and a mixture of penicillin-streptomycin (100X, 10000 U/mL) (15140, Gibco). All cells were cultured in a 37°C incubator with 5% CO_2_.

### Antibodies and chemical reagents

Primary antibodies used for western blotting include ARHGEF18 (EB06163, Everest) at 1:500, BCR (N-20, sc-885, Santa Cruz Biotechnology) at 1:1000, Cdc42 (610929, BD Transduction) at 1:1000, GAPDH (FL-335, sc-25778, Santa Cruz Biotechnology) at 1:1000, Intersectin 2 (H00050618-A01, Abnova) at 1:1000, LARG (N-14, sc-15439, Santa Cruz Biotechnology) at 1:1000, α-PIX (4573S, Cell Signaling) at 1:1000, β-PIX (07-1450, Millipore-Chemicon) at 1:2000, Rac1 (23A8, Abcam) at 1:2000, RhoA (sc-418, Santa Cruz Biotechnology) at 1:500, SOS1 (C-23, sc-256, Santa Cruz Biotechnology) at 1:1000, SOS2 (C-19, sc-258, Santa Cruz Biotechnology), Tiam1 (C-16, sc-872, Santa Cruz Biotechnology), α-tubulin (MCA77S, Serotec) at 1:2000. Secondary polyclonal antibodies conjugated with HRP for western blot were from Dako and used at 1:5000. Primary antibodies used for immunofluorescence include E-cadherin (13-1900, Invitrogen) at 1:100, ZO-1 (61-7300, Invitrogen) at 1:100. Secondary antibodies conjugated with Alexa 488 or Alexa 568 (Invitrogen) were used at 1:400. Other reagents used include Hoechst 33342 (Sigma) at 1 μg/ml, Y27632 (stock in H2O) (HA139, Sigma) at indicated concentrations, Blebbistatin (stock in DMSO) (203391, Calbiochem) at indicated concentrations, ERKi (stock in DMSO) (SCH772984, provided by Neal Rosen Lab, MSKCC) at 1μM, GSK1120212 (stock in DMSO) (S2673, Selleckchem) at 500 nM, PD0325901 (stock in DMSO) (S1036, Sellleckchem) at 500 nM, Rhosin (stock in DMSO) (EMD Millipore) at indicated concentrations, SMIFH2 (stock in DMSO) (EMD Millipore) at indicated concentrations.

### shRNAs

An shRNA library was constructed in pSUPERpuro, containing at least 3 hairpins per gene for 80 predicted human Rho GEFs. An empty pSUPERpuro vector was used as a negative control (termed ‘Control’ in the figures and ‘pSuper’ in Table 5). ARHGEF3, Rac1, RhoA, RhoC and TRIO shRNAs were obtained from the TRC library collection, in the pLKO.1 vector (MSKCC RNAi core facility) and a pLKO. 1 vector with a non-targeting sequence was used as a negative control (Sigma cat# SHC002) for these experiments. The RhoA, Rac1, Cdc42 and RhoC hairpins reported in the study were: Cdc42 sh1 (GGAGAACCATATACTCTTG), Cdc42 sh2 (GATGACCCCTCTACTATTG), Rac1 sh1 (TRCN0000004871) (TTAAGAACACATCTGTTTGCG), Rac1 sh2 (TRCN0000004873) (TAATTGTCAAAGACAGTAGGG), RhoA sh1 (TRCN0000047710) (GTACATGGAGTGTTCAGCAAA), RhoA sh2 (TRCN0000047711) (CGATGTTATACTGATGTGTTT), RhoC sh1 (CTACTGTCTTTGAGAACTATA), RhoC sh2 (GCGAACCGGATCAGTGCCTTT), RhoC sh3 (TGATGTCATCCTCATGTGCTT), RhoC sh4 (GAATAAGAAGGACCTGAGGCA). Hairpin sequences for the Rho GEFs are documented in Tables 2-3. Hairpins with un-measureable knockdown were defined as ‘off-target’ controls and were used to assess the extent of off-target effects and to define thresholds for hit-identification in the screen that minimize false detection rates (Fig. 3D).

### Virus production and infection

For virus production, 90% confluent HEK293T cells cultured in 6-well plate were transfected with lenti- or retroviral constructs using Lipofectamine 2000 (11668, Invitrogen) and Opti-MEM (31985, Invitrogen), according to the manufacturer’s instructions. The culture media were removed the day after transfection, and media were collected three times at 24 hours intervals. To infect cells, 2x10^5^ 16HBE cells were seeded in each well of a 6-well plate, and incubated on the following day with 1.5 mL virus-containing media supplemented with 1.5 μL polybrene (8 μg/μL stock, Sigma). Spin infection was performed at 2250 rpm for 30 minutes. Cells were selected starting two days after infection with 1.5 μg/mL puromycin (Sigma). Pooled selected cell lines were used for all experiments.

### Primers for PCR and qRT-PCR

Primers were designed using NCBI Primer-Blast to target all transcription variants of the gene, and to span exon-exon boundaries to avoid amplifying genomic DNA. For qRT-PCR, primers were selected to amplify a PCR product between 70-150 base pairs in length and have melting temperatures between 57°C to 63°C. Two sets of primers were examined for each gene, and the primer pair with the highest efficiency by qRT-PCR was selected for quantifying GEF expression. For some genes QuantiTect primer assays (Qiagen) were used. All primers used to quantify GEF expression are shown in Table 1.

### RNA extraction and PCR

Total RNA was extracted using the RNeasy Plus Mini Kit (74134, QIAGEN). cDNAs were prepared using Oligo dT or Random hexamer primers (IDT technology). Briefly, RNA mixtures were heated at 65°C for 5min, then chilled on ice, and mixed with 5X Reaction buffer (Thermo Scientific), RiboLock RNase inhibitor (EO0381, Thermoscientific), dNTPs (Sigma), and RevertAid reverse transcriptase (EO0441, Thermo Scientific). Reverse transcription reactions were performed using the following PCR program: 25°C for 10 mins, 42°C for 60 mins, 72°C for 10 min, then cooled at 4°C.

To examine gene expression, each PCR reaction was carried out in a total 20 μL reaction, with 100 ng cDNA as template, 0.5 μM forward and reverse primer, 2 μL 10X PCR buffer (without magnesium, Invitrogen), 1 μM dNTPs (Sigma), 1.5 mM MgCl_2_, 0.2 μL Taq polymerase (10342, Invitrogen), and 9.2 μL H2O. The PCR program used was 94°C for 3 mins, followed by 30 cycles of [94°C for 45 secs, 60°C for 30 secs, 72°C for 1 min], then followed by 72°C for 10 min and 4°C. PCR products were electrophoresed on a 2% TAE gel and stained with ethidium bromide. For quantitative Real-Time PCR (qRT-PCR), 1 μg cDNA was used per reaction, in 25 μL reactions containing 1.25 μL of 5 μM forward and reverse primers, and 12.5 μL Maxima SYBR Green/ROX qPCR Master Mix (2X) (K0221, Thermo Scientific). qRT-PCR reactions were performed on a Biorad iQ5 Multicolor RT-PCR Detection System with the following conditions: 95°C for 10 mins, 40 cycles of [95°C for 15 sec, 60°C for 60 sec] for gene expression detection, followed by 71 cycles of [60°C for 30 sec, with increase of 0.5°C per cycle] for melting curve detection. Gene expressions were normalized by the expression of GAPDH and HPRT and triplicate measurements were used for each sample.

### Western blotting

Cells were washed in ice-cold PBS and lysed in ice-cold RIPA buffer (50mM Tris-HCl pH7.4, 150 mM NaCl, 2 mM EDTA, 1% NP-40, 0.1% SDS) supplemented with 5 mM Na_3_VO_4_, 10 mM NaF, 25 mM β-glycerolphoaphate, and 1 mM PMSF. Lysates were collected by cell scraping and cleared by centrifugation at 13,000 rpm for 1 min at 4°C, and boiled in sample buffer (final concentration: 50 mM Tris HCl pH 6.8, 2% SDS, 10% glycerol, 0.1% bromophenol blue,100mM DTT) for 5 minutes. Protein concentrations were determined by BCA assay (23225, Pierce™ BCA protein assay kit, ThermoFisher Scientific). SDS-PAGE on 3-8% Tris-Acetate gels, or 4-12% Bis-Tris gels, or 12% Bis-Tris gels (NuPAGE, ThermoFisher Scientific) transfer to PVDF membranes (0.45 mm pore size, Millipore), blocking, antibody binding and ECL-mediated detection were performed as described (Durgan et al., 2014).

### Validation of expression levels and knockdown efficiencies

Western blots were used to assess protein expression levels for 9 GEFs that had validated antibodies available. qRT-PCR was applied for the remaining 71 GEFs to assess gene expression levels. The knockdown of 11 GEFs could not be assessed by qRT-PCR due to failure in production of efficient primers.

### Immunofluorescence cell-cell junctions

16HBE cells grown on glass coverslips were washed with PBS and fixed in 3.7% (v/v) formaldehyde (Sigma) in PBS for 10 minutes at room temperature. For immunostaining, coverslips were washed 3 times in PBS and blocked with BTPA buffer (0.5% BSA, 0.02% sodium azide, 0.25% Triton X-100 in PBS) for 30 minutes. Following blocking, coverslips were incubated with primary antibodies against ZO-1 and E-Cadherin (diluted in BTPA buffer) for 1 hour at room temperate and washed three times in PBS for 5 minutes. Coverslips were then incubated with secondary antibodies and Hoechst (diluted in BTPA buffer) for 1 hour at room temperature, washed 3 times in PBS for 5 minutes and mounted onto microscope slides (Fisher Scientific) using fluorescent mounting media (DakoCytomation). Epifluorescence images were acquired with an upright Imager.A1 microscope (Zeiss), equipped with an EC-Plan-NEOFLUAR 40x/0.75 objective and a Hamamatsu Orca-ER 1394 C4742-80 camera, controlled by Axiovision software (Zeiss). Scale bars were added using ImageJ.

### Wound healing assay

For wound healing assays, 3x10^6^ 16HBE cells were seeded in each well of a 6-well tissue culture plate and incubated for two days prior to wounding. Wounding was performed by scratching with a P1000 pipette tip on the confluent monolayers, in the middle of each well, and a cell scraper was used to remove half of the cells from the plate. After washing with PBS several times to remove cell debris and adding fresh 16HBE media, plates were imaged on an inverted Axiovert 200M microscope (Zeiss) equipped with an EC Plan-NEOFLUAR 10x/0.3 Ph1 objective, a Hamamatsu Orca-ER 1394 C4742-80 camera, a 37°C incubator and a CO2 controller, controlled by Axiovision software (Zeiss). The time-lapse image sequences were recorded for 16 hours at five minute intervals.

### Velocity measurements

Velocity fields were computed using custom cross correlation-based particle image velocimetry using nonoverlapping image patches of size 15 × 15 *μ*m (Zaritsky et al., 2012). The frame-to-frame displacement of each patch was defined by the maximal cross-correlation of a given patch with the subsequent image in the time lapse image sequence. The search radius was constrained to an instantaneous speed of 90 *μm* h^−1^. At a frame rate of 5 min this search radius corresponds to half the side length of a 15 × 15 *μ*m patch.

### Segmentation of monolayer contours

Monolayer contours were calculated with a segmentation algorithm that classified the image regions as either ‘cellular foreground’ or ‘background’. Our goal was to optimize robustness of the segmentation to enable unsupervised analysis of thousands of movies, each containing tens of time points. Small inaccuracies in the segmentation have little effect on the resulting kymograph, every movie was manually validated and less than 1% of the well-locations were discarded on the grounds of failed segmentation. To achieve robustness we introduced several priors to the algorithm: (1) the image contains one continuous region of ‘cellular foreground’ and one continuous region of ‘background’, (2) the contour advances monotonically over time. These priors allowed us to estimate the initial contour at time 0 and then use the segmentation at time *t* as a seed to expand the ‘cellular foreground’ to time *t*+1. The only pixels in question are those labeled as ‘background’ at time *t* and are close enough to the ‘cellular foreground’ region. The proximity threshold is calculated based on the maximal cell velocity of 90 *μm* h^−1^.

The segmentation was performed in super-pixels with a size equivalent to a 15 × 15 *μ*m patch. Following cross-correlation-based particle image velocimetry, which assigns to each super-pixel a displacement vector and a cross-correlation score defining the match of the corresponding patch signals between consecutive frames, we took advantage of the observation that super-pixels associated with ‘cellular foreground’, especially those at the front of the monolayer, had lower cross-correlation scores because of their textured and dynamic nature. We thus assigned to each super-pixel a pseudo-intensity value (1 – cross-correlation score) and applied the Rosin thresholding method (Rosin, 2001) to label the super-pixels as ‘cellular foreground’ vs ‘background’. Background super-pixels enclosed by foreground regions were re-labeled as ‘cellular foreground’ and isolated ‘cellular foreground’ super-pixels, usually attributed to debris or textured plate patterns, were re-labeled as ‘background’. We next calculated the union of the ‘cellular foreground’ region in the previous frame with the new ‘cellular foreground’ region. Morphological closing and hole-filling defined the final segmentation. For the first image in the time-lapse sequence, we defined the ‘cellular foreground’ as the union of the cross-correlation-based segmentation and a texture-based segmentation. The latter was implemented by first representing each super-pixel by the distribution of its Local Binary Patterns, a widely used representation of local texture in images (Ojala et al., 2002), followed by unsupervised K-means clustering with K = 2. This heuristic was found to robustly complement the cross-correlation-based segmentation by identifying super-pixels mislabeled as ‘background’ at larger distances from the wound edge, where cells do not sufficiently migrate and change appearance to generate a low enough cross-correlation score.

### Correction for microscope re-positioning error

During the multi-location filming, the microscope stage exhibited re-positioning errors that caused systematic shifts in all vector fields. We estimated the stage shift for every image in a time lapse sequence and corrected the vector fields accordingly. To accomplish this we exploited patches in the background region that are expected to stay in the same position and subtracted their robust mean velocity from the vector fields in the ‘cellular foreground’.

### Kymographs

Speed kymographs were constructed by calculating the average speed of all patches in spatial bands of 15 *μm* from the monolayer’s front through time. Cell directionality is defined as the absolute ratio between the velocity component perpendicular to the monolayer edge and the velocity component parallel to the monolayer edge. Each bin in the directionality kymograph was calculated as the ratio obtained by the two-component decomposition of the speed kymograph to a component normal and parallel to the monolayer front. These components were calculated by considering the orientation of the wound edge. Coordination (or short-range communication) was measured by detecting clusters of cells moving in coordination. Coordination kymographs were constructed by recording the fraction of patches that participated in these coordinated clusters for every spatial band and time-frame. Explicit detection of coordinated clusters was performed for every time-frame by applying region-growing spatial-clustering of the image-patch grid based on the correlation of their velocity fields, as described in (Zaritsky et al., 2014). Briefly, region-growing segmentation (Nock and Nielsen, 2004) started with regions containing a single image patch and iteratively merging spatially adjacent patches based on their velocity vector similarity. Two regions are merged if their similarity is lower than a given threshold and the merged region vector is updated to be the average of all contributing patch vectors. Merging is performed in ascending order of the similarity between the adjacent patches. Kymographs were calculated for cells located up to 180 μm from the monolayer and for the first 200 minutes after wound infliction. We chose 180 μm because the cell monolayers captured in the field of view of the vast majority of experiments were equal or wider than this value. We chose 200 minutes to focus our analysis on the transient phase between wound infliction and steady state collective migration. Extending the time window to 400 minutes did not change the conclusions drawn from kymograph analysis.

### Calculation of Principal Components

To obtain a compact representation of the kymographs we averaged the kymographs in 3×4 bins at a resolution of 60 μm and 50 minutes (Zaritsky et al., 2012). To further reduce this 12- dimentional feature vector we applied principal component analysis (PCA) (Jolliffe, 2002). The PCs are ranked by the spread of the data they capture. Mathematically, this is equivalent to the eigenvalues of the data covariance matrix. We used the 402 control experiments of the screen to calculate the PCA transformation for speed, directionality and coordination, and then applied the same transformation on control and knockdown experiments. First, the features were normalized (Fig. 2B) to *x*’ = (x − *μ*) / *σ*, with feature *x*, *μ* - the mean and *σ –* the standard deviation of the set of control experiments. The PCA transformation was calculated and applied to normalized features. Nearly identical PCA transformations were found when including the knockdown experiments (and for non-normalized features), indicating that the functional fluctuations between control and knockdown experiments are small compared to the day-to-day variation of control experiments. The coefficients of each PC defined the projection of the 12-dimensional feature vector into the direction of t the PC. The coefficients of PC1 were similar for all 12 features, implying that PC1 encodes the mean of speed, directionality or coordination (dependent on the considered kymograph) across time and space. The coefficients of PC2 and PC3 reflected the pattern of a temporal (PC2) or spatial (PC3) gradient (Figs. 2D, S1F, S3D; see text). It should be noted that conditions with overall reduced motility such as with CDC42 and RAC1 knockdown reduce not only PC1, but inherently flatten the spatial and temporal gradients in speed, and thus reduces PC3, but increases PC2 of this measure. This increase of PC2 is due to the positive coefficients for the (slower) speed at the onset of migration, which is minimally reduced under these perturbations, while the negative coefficients for reduced speed of perturbed experiment at later times, increases overall values. In Figures 2E, 4F-G, 5C, 5E, S1H, S2H, S3F-G, S3K we report the shifts induced by knockdown by subtraction of the PC of control experiments from the PC of the knockdown experiment. For PC1, negative values mean reduction in speed, directionality, or coordination. For PC3, which reflects the spatial gradient,negative difference values imply more immediate front-to-back propagation and faster establishment of the steady state, while for PC2 this is reversed.

### Scoring a knockdown well in relation to its daily control well

Time-lapse image sequences were acquired in at least 3 different locations in every knockdown and *in every* control well. To identify hits in the screen by comparison of knockdown and control conditions we used metrics that take into account the variability within a well and between wells. Specifically, we applied 3 metrics: i) Davies–Bouldin index (Davies and Bouldin, 1979), 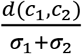, which divides the difference *d*(*c*_1_*, c*_2_) between the per-well mean values of a measurement extracted from multiple locations in the knockdown and control wells by the sum of the average distances σ_*i*_ between the measurement from individual locations in the same well and the corresponding well mean value; ii) Dunn index (Dunn†, 1974), 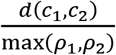, which divides the difference between per-well mean values by the maximal distance in any one well between the measurement from individual locations and the corresponding well mean value; iii) Silhouette coefficient (Rousseeuw, 1987), 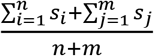, where n, m are the number of locations in control and knockdown wells, and 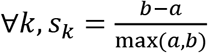, where *a* is the average difference between the measurement extracted in location *k* and the measurements extracted in all other locations of the same well, and *b* is the average difference between the measurement in location *k* and the measurements extracted in all locations of the other well.

Off-target control experiments (see text) were employed to normalize the three metrics across different PC-measurements and to define a *z-score* = (*x* - *μ*) / *σ*, with *μ* and *σ* denoting the mean and standard deviation of a particular PC metrics in the off-target control population and *x* denoting the metrics of that PC for a knockdown experiment. We defined a single combined z-score scalar for every PC-measure by accumulating the three metric-specific z-scores.

### Selecting threshold for identification of hits

The 24 off-target and 18 positive controls were used to calculate a threshold for hit identification in the screen, which has the desired property of minimizing the number of false-alarms to allow us to focus on real hits for follow-up experiments. Because the positive controls (CDC42, RAC1 and *β*-PIX) were selected based on their known effect in reducing motility, the threshold was calculated to minimize false-alarms in wound healing rate. Calculating z-scores by standardization relative to the off-target controls made the threshold transferrable to the other PC-measures. Specifically, we calculated the z-score threshold that maximizes the F-measure,defined as 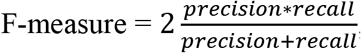,with recall 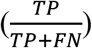 and precision 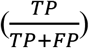derived from the true positive and false negative *FN* and false positive *FP* rates. The *TP* and *FP* rates were estimated based in on the positive controls, whereas the FN rate was were estimated based on the negative controls. Due to the small number of control experiments the threshold had low confidence. To resolve this, we applied bootstrapping to estimate the distribution of the maximal F-measure thresholds. Off-target and positive controls were randomly selected with repetition from the original groups. The process was repeated 10000 times to define the distribution of z-score values maximizing the F-measure and the final threshold of z-score = 9.8, was selected with 99.9% confidence from this distribution. There were no false-positive occurrences in 5 out of 6 PC-measures and 1 false-positive occurred for the 6^th^ PC-measure (Directionality PC2), False negative rates were 44.4%.

### Analysis of follow-up experiments

To validate the hits that were identified in the screen we replicated experiments and statistically confirmed the phenotypes. For every experiment we subtracted the mean PC-measure of the daily control locations from the corresponding PC-measure in every knockdown location. Thus each data point represents the deviation of a location to the mean control. The null hypothesis was that the data come from a distribution whose median is zero. Statistical significance was inferred using the non-parametric Wilcoxon signed rank test, and a p-value of 0.01 was selected as the significance threshold. At least 2 different hairpins were required to validate a hit. SOS1 experiments were replicated with N (number of independent experiments) = 3-6 and n (total number of locations in knockdown wells) = 18-26; RHOA-GEFs were validated with N = 6-9 and n = 23-53 and contractility experiments (treated for 48-hours) with N = 3 and n = 18. RhoGTPAses were validated with N = 6-9 and n = 24-47, *β*-PIX with N = 3, n = 11.

### Calculating similarities between different conditions

To calculate overall similarities between different experimental conditions we included the 3 first PCs for speed, directionality and coordination in a 9-dimentional vector for each well location of a knockdown experiment. As in the follow-up analysis, PC-measures in the locations of a knockdown well were related to the mean of the corresponding PC-measures in all locations of the daily control well. An experimental condition was represented as the average 9-dimensional vector of all locations in all wells under the same treatment regime. This defined a matrix with 9 rows–one per measure, and k columns, where k is the number of different conditions examined for the analysis (e.g., 8 for the analysis described in Fig. 5B). Each of the 9 measures was standardized across conditions by subtracting the mean and dividing by the standard deviation (z-scores) for each row in the matrix. This encoded the divergence of every specific condition from the mean across the examined experimental conditions. Similarity was calculated between every pair of conditions (columns) in 9-dimensions of the standardized measures using the |L1| distance metric. High values reflect dissimilarity while low values represent more similar phenotypes in this 9-dimensional space.

### Data and source-code availability

All data are made available as a public resource for the further study of GEF roles in regulating collective cell migration:

- Primers used to quantify GEF expression are in Table 1.
- shRNA hairpin sequences, knockdown efficiency (Western Blots / qRT-PCR) are in Table 2 and summarized in Table 3.
- Details of all screen experiments are in Table 4 and of all screen + follow-up studies in Table 5: molecular perturbation, raw-data file name, file format (.zvi / .tif), physical pixel size in μm, measured KD efficiency (%), experiment date.
- Screen z-scores are in Table 6, coordination z-scores from follow-up experiments are in Table 7.

Raw imaging data, processed kymographs (per location and average per day) and quantification of phenotypes for genes that were followed-up will be uploaded to a public repository upon publication.

The Matlab source code is available to the public at https://github.com/DanuserLab/MonolaverKymographs. It enables calculation of the velocity fields, segmentation of the monolayer front and the wound healing rate for every time frame in the raw time-lapse images and generates kymographs for speed, directionality and coordination for a full video. It will also produces the 12-dimentional feature vector representation and PC projection based on the transformation that was calculated for this study. We also provide a script to perform PCA and compute the transformation given a set of high-dimensional experiments. Input: M = d × n matrix. d–number of dimensions (in our case d = 12), n – number of experiments. Output: PCA transformation (wrapper for Matlab’s PCA code). This code includes means of visualization of the PCA weights (such as in Fig. 2D) to enable interpretation. Documentation and test example are included with the source code.

## Acknowledgements

We thank Andrew Jamieson for helping to package the source code and creating the repository. We thank Claudia Schaefer and Nawal Bendris for fruitful discussions and advice. This work was supported by NIH P01 GM103723 (to GD, AH, and MO). MO was also supported by NIH P30 CA008748. We dedicate this work to our mentor and colleague Alan Hall, who has initiated this project.

## Author Contribution

AH and GD conceived the study. MO, AH and GD guided AZ, YYT, MAR, SK. AZ, YYT and MAR designed the experiments. YYT, MAR and SK performed all experiments. AZ developed analytic tools and analyzed the data. AZ and YYT interpreted the data. AZ drafted the manuscript. All authors wrote and edited the manuscript and approved of its content.

## Abbreviations List

GEF: guanine nucleotide exchange factors
GDI: guanine dissociation inhibitors
16HBE: human bronchial epithelial cells from the 16HBE14o line
PCA: principal component analysis
PC: principal components
shRNA: small hairpin RNA
KD: knockdown
FBS: fetal bovine serum
PCR: polymerase chain reaction
qRT-PCR: quantitative real time PCR
ROI: region of interest

## Tables

Table 1: Primers used to quantify GEF expression for 80/81 GEFs (some had 2 different primers).

Table 2: shRNA hairpin sequences and knockdown efficiency. KD efficiency was assessed via Western Blots / qPCR. N/A indicates that the primer could not be validated by qRT-PCR due to failures in the production of efficient primers (11 GEFs). 5 GEFs were not expressed in 16HBE cells. Red font indicate (16) GEFs were all haripins had KD efficiency below 50%.

Table 3: Summary of Table 2. List of GEFs with 0, 1, 2 or 3 hairpins with KD efficiency > 50%.

Table 4: Details of all screen experiments. Details of every time-lapse experiment recorded in the screen. Every entry describes a location in a well. Each day contains a control well and multiple wells with a GEF KD. The information for every video includes the molecular perturbation in the format GENE_HAIRPIN, file name of raw data, files format (.zvi / .tif), physical pixel size in μm, measured KD efficiency (%), experiment date.

Table 6: Screen z-scores. For every combination of GEF, hairpin and experiment day the table reports the z-score for the wound healing rate and PCs1-3 for speed and directionality. Z-scores are calculated as described in Methods.

Table 7: Coordination z-scores for every combination of GEF, hairpin and experiment day.

---

**Figure S1:**
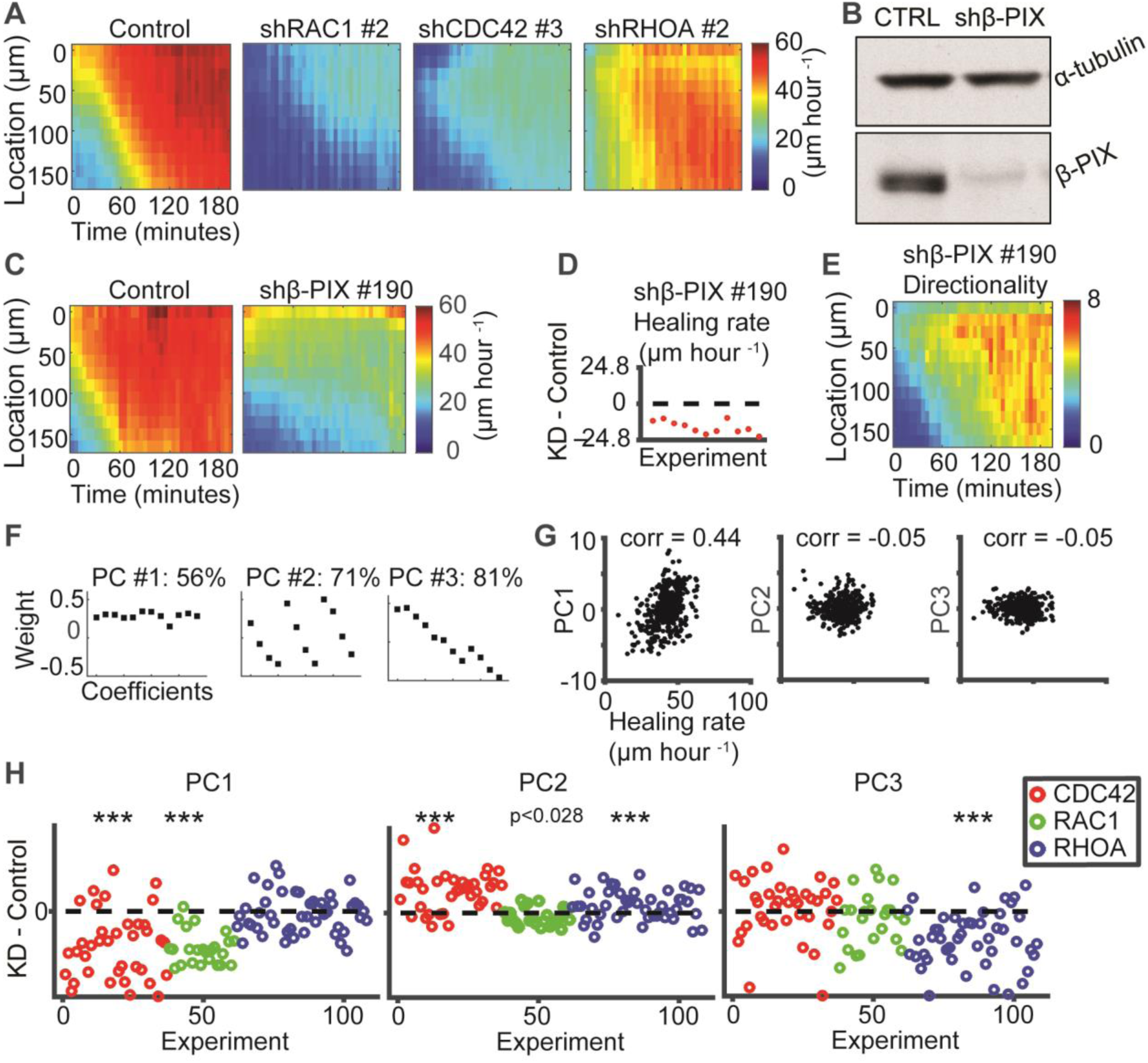
Alterations of collective motility induced by depletion of Rho GTPases or the GEF *β*-PIX. (A) Validation of Rho GTPase knockdown effect on speed kymograph by the 2^nd^ hairpin. Kymographs are averages of 3-4 locations in a well. (B) Western blot showing effective depletion of *β*-PIX. (C) Speed kymographs. (D) Wound healing rate. Each point was calculated as the difference between the migration rate in one location and the mean of the same day’s control experiment. N = 3 days, n = 11 locations, p = 0.001 via Wilcoxon signed rank test. (E) Directionality kymograph, averaged over 4 locations in one well. (F) Coefficients of the first three principal components (PCs) in directionality variation, calculated from 402 control experiments. Together, three first PCs account for 81% of directionality variation. (G) Association of PCs in directionality with wound healing rate. (H) Effect of Cdc42, Rac1 and RhoA depletion on PCs in directionality. For each experiment the difference between knockdown and control in the PC space is shown. Statistics via Wilcoxon signed rank test: p ≤ 0.01: *, p ≤ 0.001: **, p ≤ 0.0001: ***.

**Figure S2:**
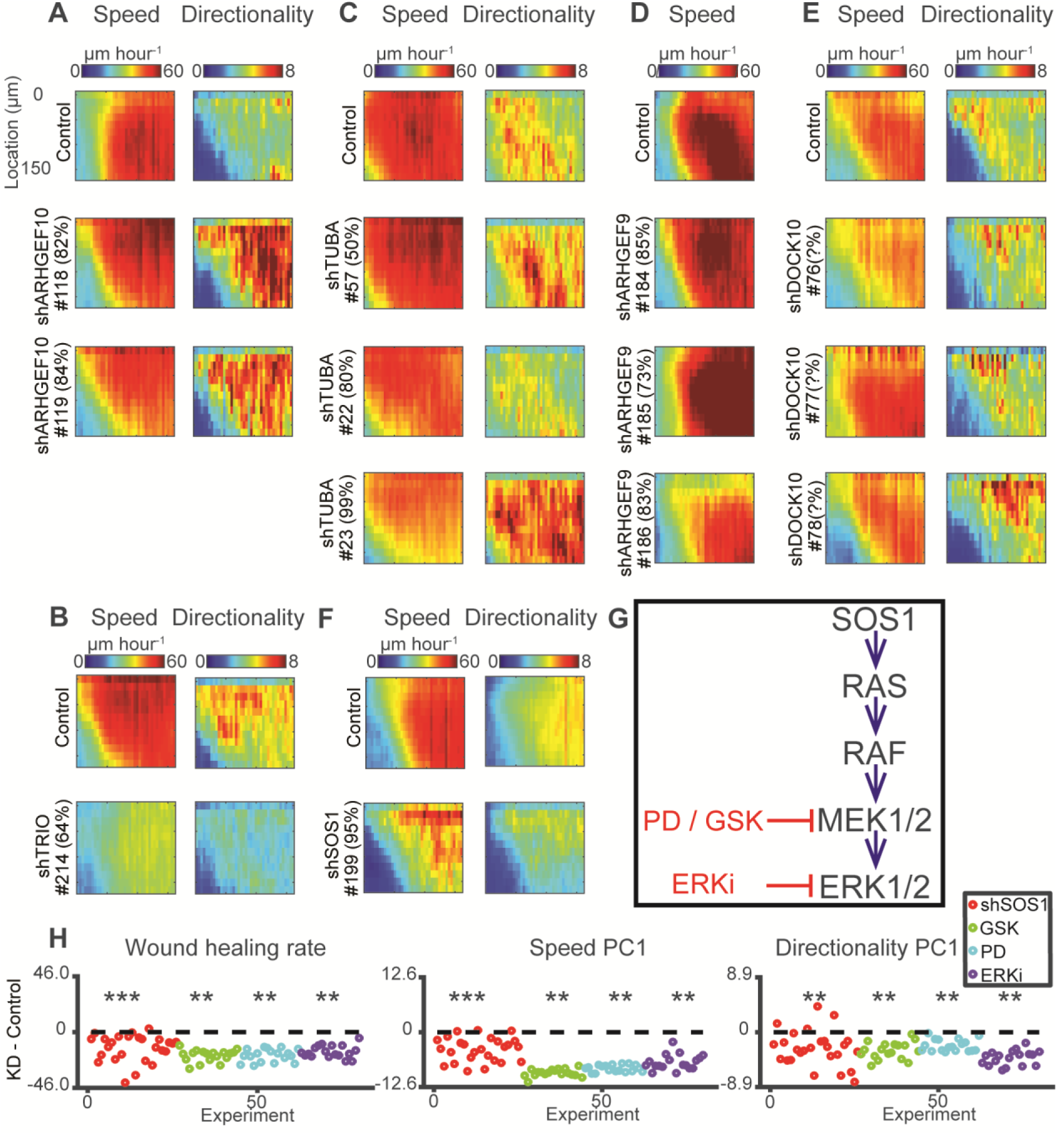
(A-E) Screen hits that were not followed-up. (A) ARHGEF10 (also named Gef10) is a RHOA-GEF (Aoki et al., 2009), showed increased directionality for two distinct validated hairpins. Was not followed-up because it was not a paralog of the other 4 RhoA-GEFs hits. (B-C) TRIO (B) is a dual GEF activating RhoA and Rac1 (Bellanger et al., 1998), TUBA (C) is a CDC42-specific GEF (Salazar et al., 2003). Both Trio and Tuba knockdowns showed reduced speed, down-regulation of Trio reduced directionality while Tuba had inconsistent directionality phenotype. Tuba showed a dose-dependency alteration: reduced speed for a more effective depletion. Both TRIO and TUBA were not followed-up because we focused on intercellular-communication alterations. (D) ARHGEF9 is a CDC42-specific GEF (Reid et al., 1999). Hairpin #184 was identified as a hit in speed PC3 - slower propagation than the control. The reason was that the control experiment on that day was an outlier - rapid front-to-back propagation in speed (PC3). Thus, the PC3 in the knockdown experiment was erroneously identified as hit. (E) DOCK10 is a dual GEF for CDC42 and RAC1 (Ruiz-Lafuente et al., 2015). Hairpin #78 came as a hit in directionality PC2. None of the 3 primers for DOCK10 worked with qRT-PCR to assess knockdown efficiency thus were labeled as 'unknown'. Together with the inability to observe a similar phenotype in the two other hairpins, and the fact that temporal gradient is not an intercellular phenotype, we decided not to follow-up on DOCK10. (F) Screening PC2, encoding temporal gradient in speed (left) and directionality (right). Grouping include positive controls, off target controls, GEFs that were screened but not identified and/or validated as hits and the validated hits (for other measures). DOCK10 was the only GEF that altered PC2 for directionality above the screening threshold. (F-H) SOS1-RAS pathway regulates collective cell migration. (F) Speed kymographs. (G) SOS-RAS pathway. Inhibition of MEK with GSK or PD and of ERK by ERKi. (H) SOS1-RAS follow-up experiments. Each data point is the subtraction of the mean measured phenotype of daily controls from the certain location within a well (KD-Control). Left-to-right: wound healing rate, PC1 speed (encoding magnitude) and PC1 directionality. N – number of wells, n – number of locations (N = 3-6, n = 18-26), 1 well per daily experiment, number of daily experiments (replicates) ≥ 3. SOS1: N = 6, n = 26, GSK, PD, ERKi: N = 3, n = 18. Statistics via Wilcoxon signed rank test: p ≤ 0.001: **, p ≤ 0.001: ***. Note that with more replicates, we were able to reveal a systematic reduction of speed and directionality that were not identified by the primary screen.

**Figure S3:**
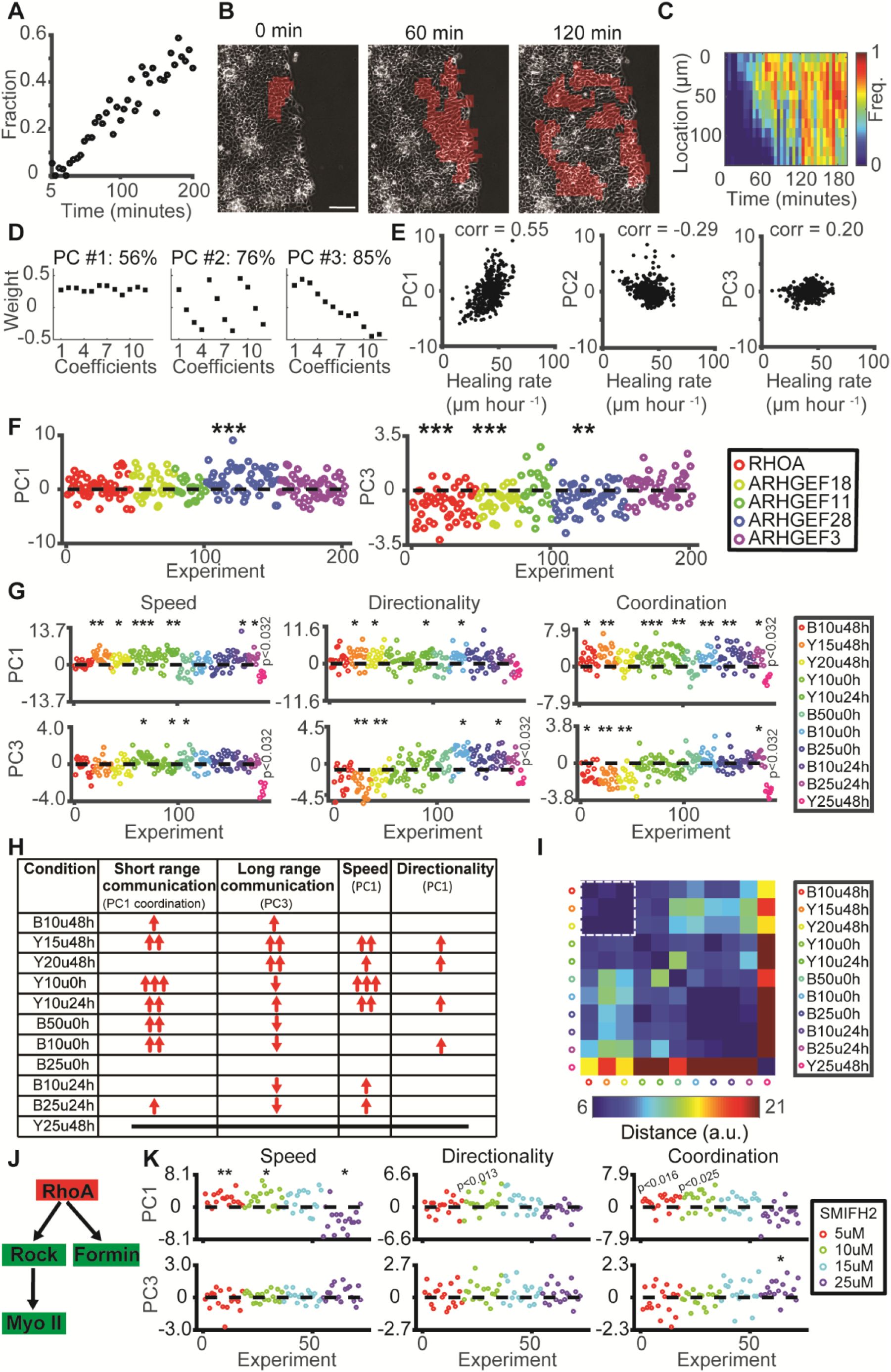
Effects of RhoA GEF depletion on short-range communication as quantified by the coordination parameter (A-F) and effects of perturbation of actomyosin contractility on intercellular communication (G-I). (A) Fraction of cells migrating in coordinated clusters (coordination) over time. (B) Representative formation of clusters in a control experiment. Scale bar = 100 μm. (C) Kymograph of coordination. (D) Coefficients of the first three Principal Components (PCs) of coordination extracted from N = 402 control experiments. (E) Association of PCs with wound healing rate. (F) Difference in PC1 and PC3 between GEF KD and control. Each data point is the subtraction of the mean PC value of the daily controls from the PC value extracted in one well location. Same experiments as in Fig. 4G. (G) Shifts in spatiotemporal dynamics of speed, directionality, and coordination upon inhibition of myosin II by pre-incubation of cells with Blebbistatin or Y27632 (denoted ‘B’ or ‘Y’) at low dosages (10, 15, 25 or 50 uM, denoted ‘u’) for 0, 24 or 48 hours (denoted 0, 24 or 48h) prior to wound healing experiment. Each point represents the difference in PC of the indicated variable between GEF KD in one well location and the mean of the daily control; N – number of wells, n – number of locations (N = 1-5, n = 6-35), 1 control well per daily experiment. B10u48h: N = 3, n = 18; Y15u48h: N = 3, n = 18; Y20u48h: N = 3, n = 18; Y10u0h: N = 5, n = 35; Y10u24h: N = 2, n = 11; B50u0h: N = 3, n = 15; B10u0h: N = 3, n = 17; B25u0h: N = 4, n = 23; B10u24h: N = 2, n = 13; B25u24h: N = 2, n = 10; Y20u48h: N = 1, n = 6. Statistics via Wilcoxon signed rank test: p ≤ 0.01: *, p ≤ 0.001: **, p ≤ 0.0001: ***. (H) Summary of alterations induced by contractility inhibition. Number of arrows corresponds to p-values in F and G. (I) Pairwise distance matrix measuring the similarity of perturbation effects (Methods). Dashed white line marks the 48-hour treatment functional cluster. (J) RHOA is upstream of the ROCK-myosin II and the mDia pathways. (K) Increased speed and subtle increase in directionality and coordination upon pan-inhibition of formin-family members by acute treatment with low dosages of SMIFH2 (5, 10, 15 or 25 uM) (Rizvi et al., 2009). Each point represents the difference in PC of the indicated variable between GEF KD in one well location and the mean of the daily control; N = 3 and n = 18 for each experimental conditions, 1 control well per daily experiment. Statistics via Wilcoxon signed rank test.

